# A generative-AI framework for target-Specific MicroRNAs towards RNAi-based drug design

**DOI:** 10.64898/2026.05.07.723585

**Authors:** Jiayao Gu, Yue Li

## Abstract

MicroRNA (miRNAs) are small non-coding RNAs that regulate gene expression by binding to the target messenger RNA (mRNA), whose versatility has inspired RNA-interference (RNAi)-based drug designs. However, off-target effects lead to unintended gene silencing and toxicity. Existing methods suffer from experimental data scarcity and fail to effectively integrate target specificity into designing de novo small interference RNAs (siRNA). To overcome the above challenges, we present SpeciMiR, a specificity-guided generative framework that synthesizes target-conditioned miRNAs. By training on a large experimental data containing 2.2M miRNA-mRNA pairs, SpeciMiR minimizes off-target effects with enhanced on-target potency. As a result, SpeciMiR-generated miRNAs bind more strongly to the target mRNAs than the observed miRNAs and much less so to off-target mRNAs. We tested SpeciMiR on mRNA targets for liver disease, for which 6 FDA-approved siRNA-based drugs were available. SpeciMiR recovers binding regions that correspond to FDA-approved siRNA drugs across 3 targets, and demonstrates greater structural specificity for on-target mRNAs than for off-target mRNAs. Together, SpeciMiR offers an AI solution to synthesize miRNA-inspired and target-specific siRNA sequences towards RNAi-based drug design. Code/data is available at https://anonymous.4open.science/r/SpeciMiR-278F

## 1 Introduction

MicroRNAs (miRNAs) are small endogenous non-coding RNAs of approximately 22 nucleotides that regulate gene expression by binding to target messenger RNAs (mRNAs), leading to their degradation or translational repression [1]. Canonical miRNA targeting is primarily mediated by near-perfect base pairing between the miRNA seed region, usually nucleotides 2-8 from the 5’ end, and complementary sites in the mRNA 3’ UTR. Each miRNA can target hundreds of mRNAs, forming post-transcriptional regulatory networks that control diverse biological processes such as cell proliferation, differentiation, and apoptosis [2]. Dysregulation of miRNA expression has been implicated in numerous diseases, including cancer and neurological disorders [2, 3].

The biology of miRNAs inspired the development of exogenous small interference RNA (siRNA) and RNAi-based drugs for human diseases [4]. Similar to miRNAs, siRNAs target mRNAs by binding to 22-26 nt segment of the target sequence through complementary base pairs and trigger downstream cleavage, degradation or translational repression [5]. This mechanism gives biologists the ability to silence any gene by designing a complementary short sequence to the target mRNA [4, 5].

However, off-target effects occurs when siRNAs bind to unintended targets because of imperfect complementarity to a short segment of the 3’ end of the target mRNA sequence, which leads to unintended gene silencing and toxicity [6, 7]. Direct experimental data on siRNA transfections are scarce. For instance, the widely used BIOPREDsi dataset [8] for most siRNA models includes only 2,431 siRNAs for 34 genes. Significant efforts have been invested into eliminating off-target effects of RNAi-based drug design for more than 2 decades, mainly in chemical modification and computational design [9]. Existing computational tools design and rank siRNA candidates with post-hoc off-target filtering [10, 11, 12]. On the other hand, endogenous miRNA-mRNA interactions can be measured by genome-wide high-throughput sequencing platforms (e.g., enhanced Cross Linking and ImmunoPrecipitation or eCLIP), leading to more than 1.25 million miRNA-mRNA pairs, providing a much bigger training data for siRNA designs [13]. Moreover, EVA [14] demonstrates that large language models can generate realistic miRNA sequences conditioned on RNA class. However, EVA generates from a family-level prior without target conditioning. GEMORNA [15] uses Transformer encoder-decoders to design functional coding RNAs for translational efficiency and stability but it is not optimized for designing target-specific siRNAs.

In this study, we introduce **SpeciMiR**, the first method to generate de novo miRNAs, conditioned on any target mRNA, with specificity optimization integrated into the training objective. Our main contributions are as follows:

- We propose a specificity-guided target-conditioned generative framework for de novo miRNA synthesis. To the best of our knowledge, this is the first generative method for de novo miRNA design targeting any specific mRNA sequence while ensuring high specificity.
- SpeciMiR leverages a pretrained discriminator to improve specificity of the generator and account for off-target effects conditioned on an mRNA input.
- Comprehensive experiments show that almost all of the SpeciMiR-generated miRNAs contain valid binding sites and demonstrated significant mitigation on off-target effects. These include recapitulating the target sites of disease genes tackled by Food Drug Administration (FDA)-approved RNAi-based drugs.

## 2 SpeciMiR Model

As an overview, SpeciMiR has two training phases: in Phase 1, we finetuned an RNA encoder and miRNA-mRNA discriminator to distinguish experimentally observed miRNA-mRNA pairs from false pairs; in Phase 2, we trained a generator by propagating the gradients from the frozen discriminator loss to generate target-specific miRNA sequences (Fig. 1). For the discriminator, SpeciMiR leverages a pretrained model, MiRformer [16], which is a dual-transformer framework trained to predict interacting miRNA-mRNA pairs (Fig. 1A). Briefly, the miRNA encoder uses full self-attention to capture global dependencies within the short miRNA sequence; the mRNA encoder adopts sliding-window self-attention to efficiently scale to long mRNA contexts and model local binding regions following the Longformer design [17]. For the generator, SpeciMiR leverages the pretrained mRNA encoder from MiRformer to extract mRNA sequence features. It then uses a transformer decoder to generate miRNAs given the target mRNA embedding as an input (Fig. 1B). The details of each component is described in the following subsections.

**Figure 1:**
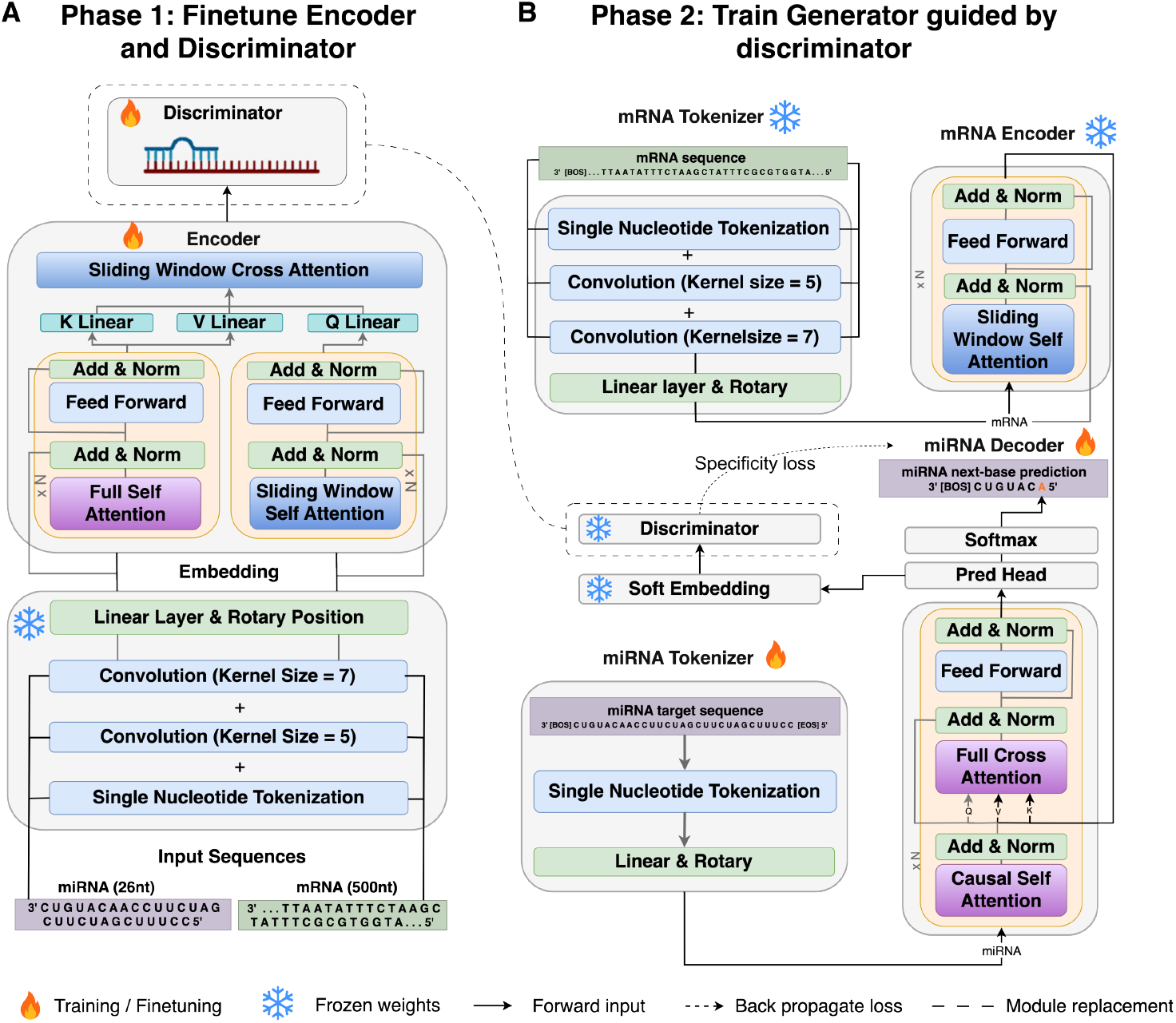
SpeciMiR overview. A) Phase 1: finetuning RNA encoder and discriminator on experimental eCLIP datasets containing 1.25M positive miRNA-mRNA pairs and 1.25M negative pairs containing binding sites. Discriminator is trained to distinguish the positive pairs from the negative ones. B) Phase 2: training the miRNA generator by guiding the updates with specificity loss from the finetuned discriminator.

### 2.1 Phase 1 - finetuning RNA encoder and discriminator on eCLIP datasets

To effectively distinguish true miRNA-mRNA interaction pairs from false seed-matched pairs, we trained a discriminator on miRBench eCLIP data, which consists of 2.2 million pairs with balanced number of positive and negative miRNA-mRNA pairs [18]. Negative pairs contain canonical binding sites but do not result in miRNA-mRNA interactions (Section 3.1).

#### Discriminator architecture

Given a miRNA sequence **m** and an mRNA sequence **x**, the discriminator processes them through separate encoders, fuses the representations via cross-attention, and produces a binding score:

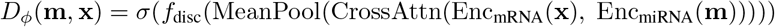

where Enc_mRNA_ and Enc_miRNA_ are Transformer encoder, with sliding-window self attention for mRNA. CrossAttn computes cross-attention with mRNA as query and miRNA as key/value, *f*_disc_ is a feed-forward binding head, and *σ* is the sigmoid function.

#### Discriminator Loss

The discriminator is trained with binary cross-entropy over a dataset of *N* miRNA–mRNA pairs with binary interaction labels *c*_*i*_ ∈ {0, 1}:

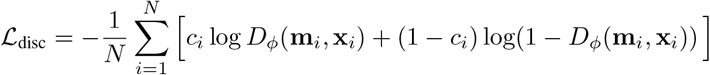

The embedding layers and both sequence encoders are frozen to preserve pretrained sequence representations. Only the cross-attention layer and discriminator are updated at different pace:

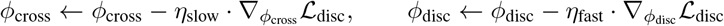

where *η*_slow_ *< η*_fast_ (by default, *η*_slow_ = 5e-5, *η*_fast_ = 1e-4), allowing the cross-attention layer to adapt gently while the prediction head adapt faster on the harder dataset (Section 3.1 Dataset). A linear warmup followed by cosine annealing is applied to both learning rates.

After Phase 1, the discriminator is frozen entirely (*ϕ* fixed) and used as a differentiable scoring function in Phase 2.

### 2.2 Phase 2 - discriminator-guided generator training with off-target risk awareness

The generator is trained with a joint loss comprising a generation quality term and a specificity term:

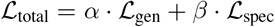

where *α* = 0.5 controls the weight of generation quality (anchoring outputs to biologically valid miRNA sequences) and *β* = 0.7 controls the weight of the specificity objective.

#### Generation loss

The generation loss is a standard autoregressive cross-entropy loss under teacher forcing. Given a target mRNA **x**_target_ and its corresponding true miRNA sequence **y** = (*y*_1_, *y*_2_, …, *y*_*T*_), the generator parameterized by *θ* is trained to maximize the likelihood of each token conditioned on all preceding tokens:

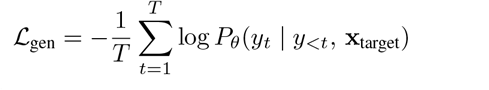

This term serves as a regularizer, preventing the generator from drifting into degenerate outputs that exploit the discriminator without producing biologically valid miRNA sequences.

#### Specificity loss

The specificity loss encourages the generated miRNA to interact strongly with the intended target mRNA while avoiding interactions with off-target mRNAs. For each training sample, we evaluate the generator’s output against the target mRNA and *K* randomly sampled negative mRNAs containing the same seed matches:

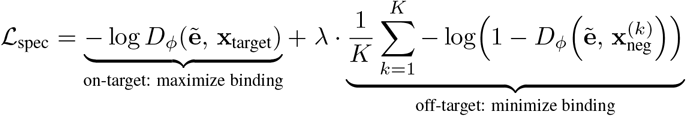

where 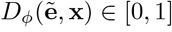 is the predicted probability of interaction between the miRNA representation 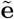 and mRNA sequence **x** by the frozen discriminator, and *λ* = 0.5 controls the penalty for off-target binding. Minimizing the first and second term pushes the discriminator’s on-target score toward 1 (strong binding) and the off-target scores toward 0 (no binding), respectively. The balance between these two objectives is controlled by *λ*.

#### Optimization details for the generator

The generator autoregressively outputs discrete tokens sampled from the nucleotide probability distribution. We need to create a differentiable path from the discriminator loss to the generator’s weights, where gradient-based optimization can be performed. To that end, we provide the soft embedding input from the generator to the discriminator.

Specifically, at each decoder position *t*, the generator produces logits **z**_*t*_ ∈ℝ^*V*^ over the nucleotide vocabulary (i.e., U, C, G, A). We convert these to a continuous embedding via temperature-scaled softmax projection: 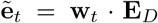 where 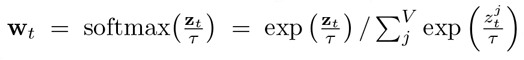, **E**_*D*_ ∈ ℝ^*V ×D*^ is the frozen discriminator’s token embedding matrix and *τ >* 0 is a temperature parameter. The resulting soft embedding 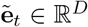 is a weighted average of all token embeddings, with weights determined by the generator’s output distribution. The temperature *τ* controls the sharpness of this distribution: as *τ* → 0, the soft embedding approaches a hard one-hot lookup; as *τ* increases, the distribution becomes more uniform and gradients flow freely but the representation becomes less precise. We use *τ* = 0.5 as a default.

The soft embedding 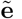 is then passed through the frozen discriminator layers (i.e., CNN tokenizer, miRNA encoder, cross-attention with encoded mRNA, and feed-forward head) to produce the interaction score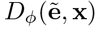. Since all operations are differentiable, gradients propagate through ℒ_spec_, the frozen discriminator to the generator’s logits **z**_*t*_ and decoder parameters *θ*. Notations and parameter values are listed in Table A.1.

## 3 Experiments

### 3.1 Experimental Setup

#### Datasets

TargetScan dataset - To train SpeciMiR on general seed binding rules, we obtained TargetScan training dataset, which contains 257,015 miRNA-mRNA pairs [19]. We constructed three datasets corresponding to three mRNA lengths: 30nt, 100nt and 500nt, each composed of 257,015 miRNA-mRNA pairs and mRNAs containing canonical seed matches. We also constructed three validation datasets corresponding to the three mRNA lengths, each containing 22,847 miRNA-mRNA pairs with canonical seed match.

To finetune SpeciMiR’s discriminator and generator on independent experimental data, we used the miRBench data [18], which provides bias-corrected miRNA binding site interaction datasets derived from CLASH and chimeric eCLIP experiments. miRBench addresses the miRNA frequency class bias in existing datasets by ensuring that miRNA family frequencies are balanced between positive and negative classes. We used the Manakov2022 dataset from miRBench containing 1.2M miRNA-mRNA with class-balanced label distribution (1:1 positive to negative ratio). Binding site sequences are short (50 nt) chimeric read fragments. We split Manakov2022 set [13] into training set (N=1,065,322) and validation set (N=187,998). We evaluated SpeciMiR on Manakov2022 test set (N=168,342 positives, chimeric eCLIP) also with 1:1 positive-to-negative ratio.

#### Evaluation of the generated miRNAs

We evaluated SpeciMiR on Manakov2022 test set by measuring its generated miRNAs sequence quality that include mean length, canonical seed match rates, and sequence diversity (i.e., the proportion of unique miRNA sequences). We scored the miRNA-mRNA pairs by the binding probability from the discriminator, energy change Δ_*G*_ by RNACofold [20], and Minimum Free Energy (MFE) of RNA alignment by RNAhybrid [21].

In addition, we performed a case study on 6 FDA-approved RNAi-based drugs. Specifically, we extracted siRNA sequences from [5] and the target mRNA sequences of the disease genes in 3’UTR or CDS regions from NCBI. To match the training mRNA lengths, we extract a 50-nt window that contains the siRNA binding sites from the full mRNA sequences. In total, we constructed 6 siRNA-mRNA pairs. Then we fed target mRNA sequences to SpeciMiR to synthesize and evaluate the generated miRNAs on binding site overlap, specificity and thermodynamics metrics. We compared the generated miRNA with the established siRNA sequences on binding site overlap measured by Intersection over Union (IoU), which is defined as the ratio between the overlap and union of the two groups: 1. base pairs between the generated miRNAs and the target mRNA, 2. base pairs between the groundtruth siRNAs and the target mRNA.

#### Two-stage SpeciMiR training

First, we pre-trained the decoder on TargetScan [19] to generate miRNAs conditioned on 30-, 100-, and 500-nt mRNA inputs containing annotated seed matches. Second, we fine-tuned the decoder on experimental eCLIP/CLASH data from Manakov2022 using a frozen discriminator trained to distinguish true miRNA–mRNA interactions from seed-matched negatives. Training details and hyperparameters are provided in Appendix A.1.

## 4 Results

### 4.1 SpeciMiR synthesizes target-specific miRNA candidates

We evaluated SpeciMiR on 30-, 100-, and 500-nt mRNA inputs. As expected, per-base miRNA prediction accuracy improves as input length decreases (Fig. A.2). To assess whether generated miRNAs preserve target complementarity, we aligned their seed regions, defined as positions 2–8 nt from the miRNA 5’ end, to the corresponding target mRNAs and quantified canonical 6mer, 7mer-a1, 7mer-m8, and 8mer matches. Across 5,712 tested 500-nt targets, 99.30% of generated miRNAs contained valid seed matches, with 45.49% containing 8mer matches (Fig. 2a; Table A.3).

**Figure 2:**
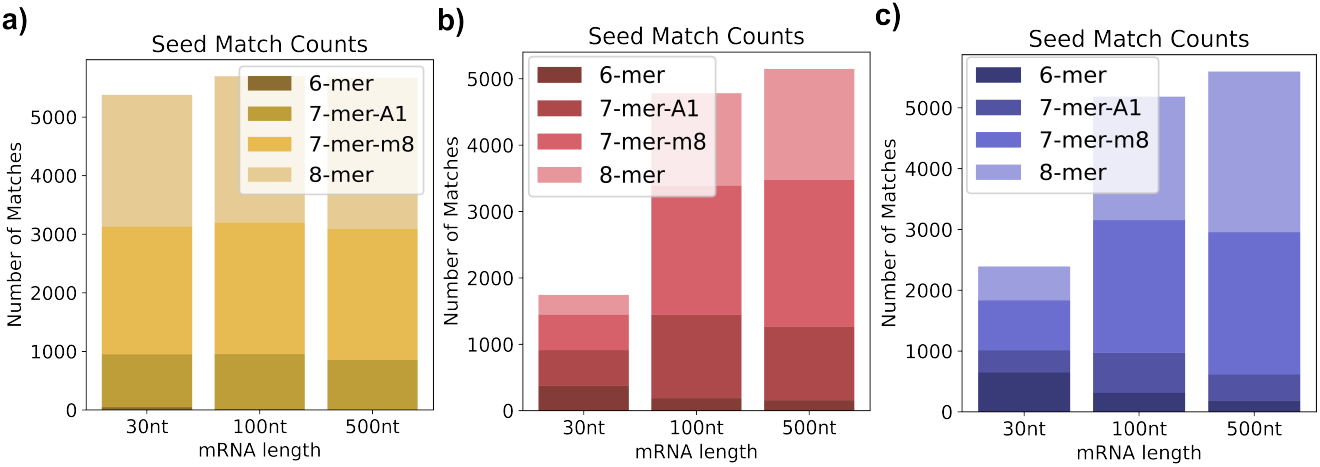
Distribution of canonical seed match types in SpeciMiR-synthesized miRNAs. Seed match categories include 6mer, 7mer-a1, 7mer-m8, and 8mer, shown as the counts of each category and their percentages relative to the total number of seed matches. a) Distribution of seed types on the original mRNA. b) Distribution of seed match types between the 2-8 nt position at the 5’ ends of the generated miRNA and the seed-perturbed mRNA. c) Distribution of seed types between any 6-8 nt segment of the miRNA and the seed-perturbed mRNA.

We further tested robustness of SpeciMiR by perturbing seed-matching regions in the target mRNAs and regenerating miRNAs. First, we used strictly the seed region - position 2-8 nt at miRNAs 5’ ends - and searched for complementary seed matches in mRNAs. As expected, perturbation substantially reduced seed-match recovery for 30-nt inputs, from 94.17% to 30.55%, but had a smaller effect on longer inputs: total seed-match rates decreased by 16% for 100-nt targets and 9% for 500-nt targets (Fig. 2b). Second, we relaxed the search by scanning the entire miRNA for any 6-8 nt segment complementary to the perturbed mRNAs. The perturbation effect was further compensated in 100-nt and 500-nt mRNA sequences, yielding a decrease by 9% and 1.35% respectively (Fig. 2c). These results show that SpeciMiR synthesizes miRNAs with valid target-complementary seed regions and can partially restore seed complementarity after target perturbation, with stronger robustness for longer target sequences. The robustness is useful when single-nucleotide polymorphism (SNPs) in the seed regions of a disease-causing gene may disrupt the binding of endogenous miRNAs causing dysregulation of the gene but can be restored via exogenous siRNAs.

### 4.2 Discriminator-guided miRNA generation with off-target-risk awareness

To allow miRNA generation to be aware of off-target risk, we finetuned discriminator on the experimental eCLIP datasets from miRBench [18]. Then we froze the discriminator’s weights and finetuned the decoder on generating miRNAs conditioned on the on-target mRNA. To minimize the off-target loss, we sampled K off-target mRNAs containing the same seed match region in their sequences (Section 2.2). We observed that by introducing the specificity loss, the binding probability on on-target mRNAs are much higher than the binding probability on off-target mRNAs (0.852 vs 0.335), and specificity gap is 0.517 (Table A.6). This shows that specificity loss correctly guided the decoder to generate miRNA that binds stronger to target mRNAs than off-target mRNAs.

### 4.3 Generated miRNA sequence properties

We evaluated SpeciMiR on the quality of the generated miRNA sequences on the Manakov2022 test set (Section 3.1 Experiments). We noticed that the sequence diversity for the generated miRNA is much higher than the observed miRNAs from the Manakov2022 test set (60.4% vs. 0.37%, Table 1). The low sequence diversity among the real miRNAs reflect the biology evolution, where only a small set of conserved miRNAs target many mRNAs [22]. In addition, the canonical seed match rate for the generated miRNAs is 34.97%, which is much higher than the observed miRNA (15.32%). This shows that SpeciMiR is able to generate miRNAs with higher canonical seed match composition than the observed miRNAs.

**Table 1:**
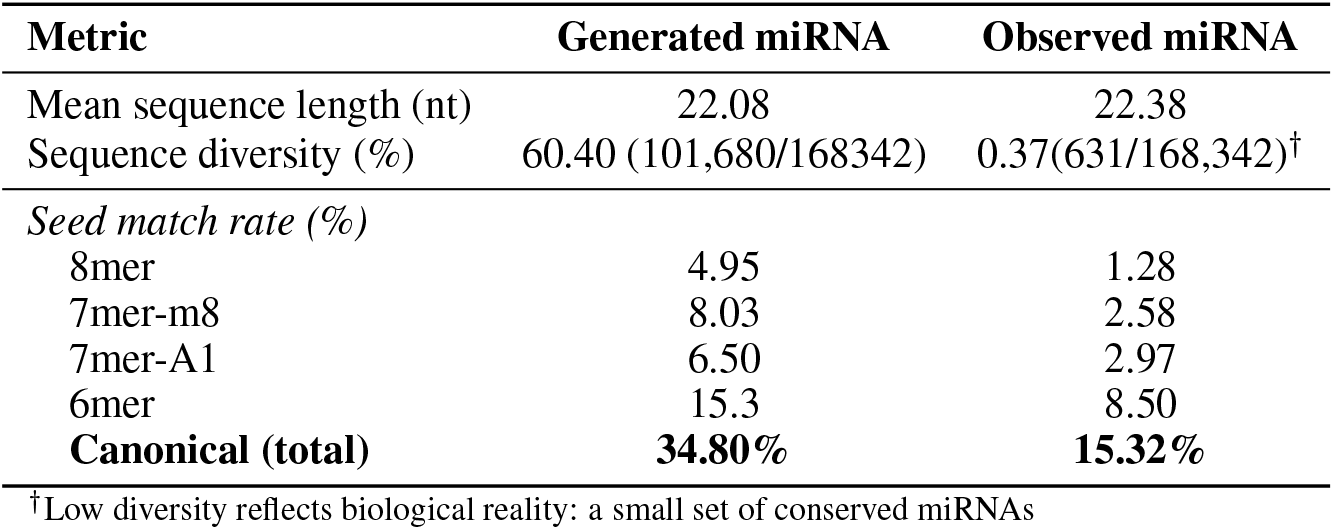
Generated sequence quality on the Manakov2022 test set (168,342 positive pairs). Seed match denotes the percentage of miRNAs exhibiting canonical seed complementarity (positions 2–8) to their paired target mRNA.

In addition, the discriminator scored the generated miRNAs with higher probabilities than the observed miRNAs (0.652 vs 0.406), which shows that the generated miRNAs are more likely to bind to the target mRNAs than the observed miRNAs (Table 2, Fig. A.3a). One caveat is that the generator is optimized for the discriminator loss, whereas the real miRNAs are under more constraints (e.g. Drosha and Dicer cleavage, AGO2 binding, etc.).

**Table 2:**
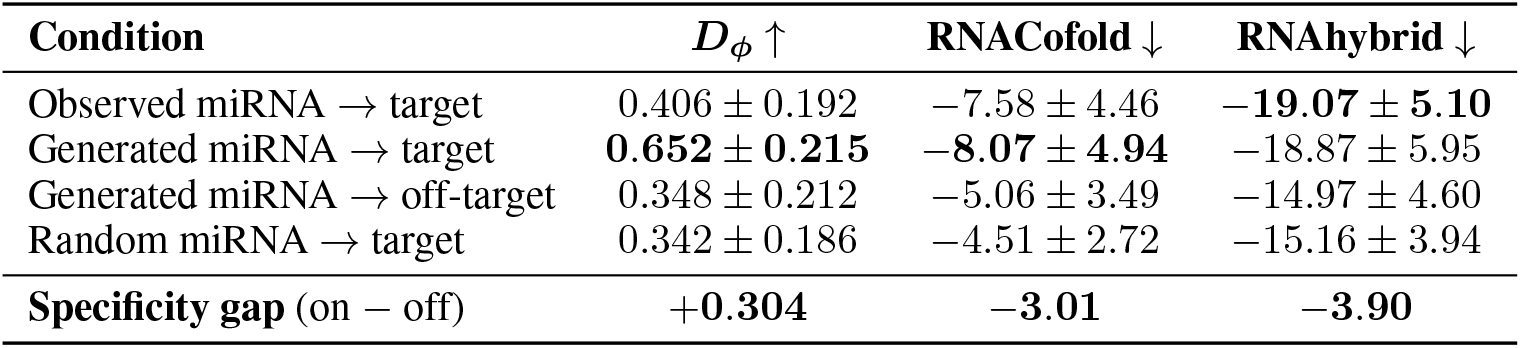
Specificity evaluation with independent scorers. **D**_*ϕ*_: finetuned discriminator (higher = stronger predicted binding). **RNACofold**: co-folding Δ*G*_binding_ (kcal/mol). **RNAhybrid**: minimum free energy (kcal/mol). More negative energies indicate stronger binding. All values reported as mean ± std over 2,000 sampled pairs (10,000 for off-target, *K* = 5 per generated miRNA).

Furthermore, RNACofold-predicted thermodynamics of the generated miRNAs are slightly more stable than the observed miRNAs (−8.07 kcal/mol vs −7.58 kcal/mol), meaning the generated small RNA forms more stable miRNA-mRNA duplexes than the observed miRNAs (Table 2, Fig. A.3b). In contrast, the generated miRNAs bind with off-target mRNAs with higher Δ*G*_binding_ than on-target mRNAs (−5.06 kcal/mol vs −8.07 kcal/mol) (Table 2).

Lastly, the RNAhybrid-predicted MFE of generated miRNAs are comparable to the real miRNAs (−18.87 kcal/mol vs −19.07 kcal/mol) (Table 2, Fig. A.3b). MFE is higher for off-target mRNAs (−14.97 kcal/mol vs −18.87 kcal/mol), meaning the generator forms less stable duplexes between the generated miRNAs and off-target mRNAs (Table 2).

As expected, all SpeciMiR-generated miRNAs bind better to on-target mRNAs than randomly generated miRNAs.

### 4.4 Ablation studies

We trained ablated variants of the full SpeciMiR to quantify the contribution of each component of the loss function. Each variant was trained on the complete Manakov2022 training set for 2 epochs and evaluated on the Manakov2022 test set (Table 3).

**Table 3:**
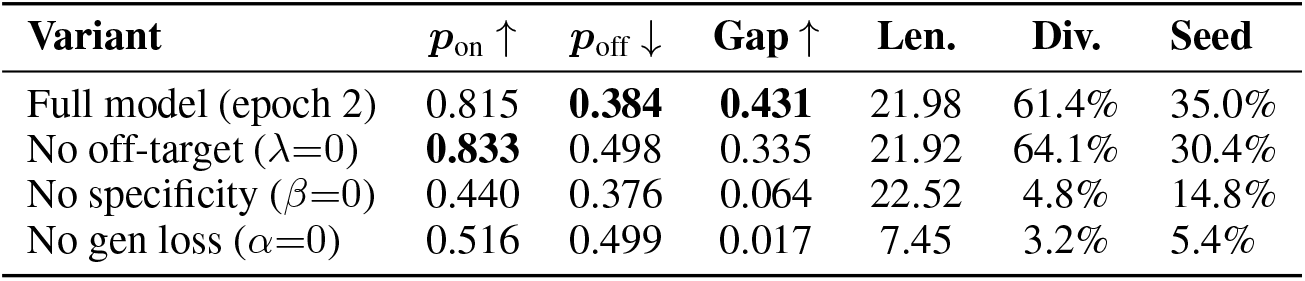
Ablation study. Full model (*α*=0.5, *β*=0.7, *λ*=0.5) trained on Manakov2022 train, evaluated on Manakov2022 test. *p*_on_: discriminator score against target mRNA. *p*_off_: discriminator score against off-target mRNA.

Removing the generation loss term (*α* = 0.0, *β* = 1.0, *λ* = 0.5) caused mode collapse, yielding very low diversity (3.2%) and short average sequence length (7.45 nt).

Removing the specificity loss (*α* = 1.0, *β* = 0.0, *λ* = 0.5) produced syntactically valid sequences but failed to discriminate between on-target and off-target mRNAs (*p*_*on*_ ≈ *p*_*off*_).

Removing the off-target loss (*α* = 0.5, *β* = 0.7, *λ* = 0.0) led to strong binding to target mRNAs (*p*_*on*_ = 0.833), but also showed high binding affinity to off-target mRNAs (*p*_*off*_ = 0.498).

Therefore, the full model achieves the best balance between sensitivity and specificity.

### 4.5 Benchmarking with miRNA design methods

We compared against two baselines for assessing generation quality: miRarchitect [23] and EVA [14] (Section A.2; Table A.7). Note that EVA is not an optimal method due to its unconditional miRNA generation. Nonethless, we used EVA [14] to generate 1000 miRNAs in total and filtered for 40 best miRNAs candidates. We used miRarchitect to generate miRNAs on 40 randomly selected mRNA targets from Manakov2022 test set. Additionally, we sampled 20 off-target mRNAs from the test set for each target mRNA that shared the same seed regions with the real miRNAs. We scored the interactions between the generated miRNAs with on and off-target mRNAs (Table A.7). Further details are described in Appendix A.3.

MiRarchitect showed larger thermodynamic specificity gaps between on-target and off-target mRNAs, with specificity gap of 23.2 kcal/mol Δ*G* and 26.4 kcal/mol MFE. This is expected because miRarchitect designs perfectly complementary miRNAs against target mRNA segments, scores and ranks the miRNA candidates in terms of GC content, secondary structure and energy, yielding more thermodynamically stable miRNAs [23]. SpeciMiR generates partially complementary miRNA-like sequences learned from *in vivo* AGO2 eCLIP-derived interactions (Section 3.1 Dataset). Importantly, while miRarchitect generates miRNAs with high target specificity, it cannot explore miRNA sequence space. This design results in failure to produce a valid miRNA for any target mRNA sequence - among the 40 given target sequences, miRarchitect returns only 19 miRNAs resulting in a response rate of 47.5%. On the other hand, SpeciMiR can generate miRNAs for any given target mRNA, yielding 100% response rate. This capability is attributed to its AI-based generative framework and efficient autoregressive training stratgey, which enable SpeciMiR to learn underlying biological principles from millions of miRNA-mRNA interaction pairs.

### 4.6 SpeciMiR-generated siRNA versus FDA-approved siRNA drugs on disease target genes

To evaluate SpeciMiR on siRNA drug designs, we use it to generate siRNAs to regulate disease target genes, for which there have been 6 FDA-approved siRNAs. To evaluate binding site overlaps, we locally aligned each generated miRNA sequence with the target mRNA sequence and compared the aligned regions with the siRNA binding sites (Fig. 3). We observed that 3 of the 6 generated miRNAs targeted the same region of the mRNAs as siRNA (IoU ≈ 0.5, with IoU on Vutrisiran reached 0.54). This shows that SpeciMiR is able to identify the most pharmaceutical siRNA binding sites entirely by learning from natural miRNA-mRNA binding rules.

**Figure 3:**
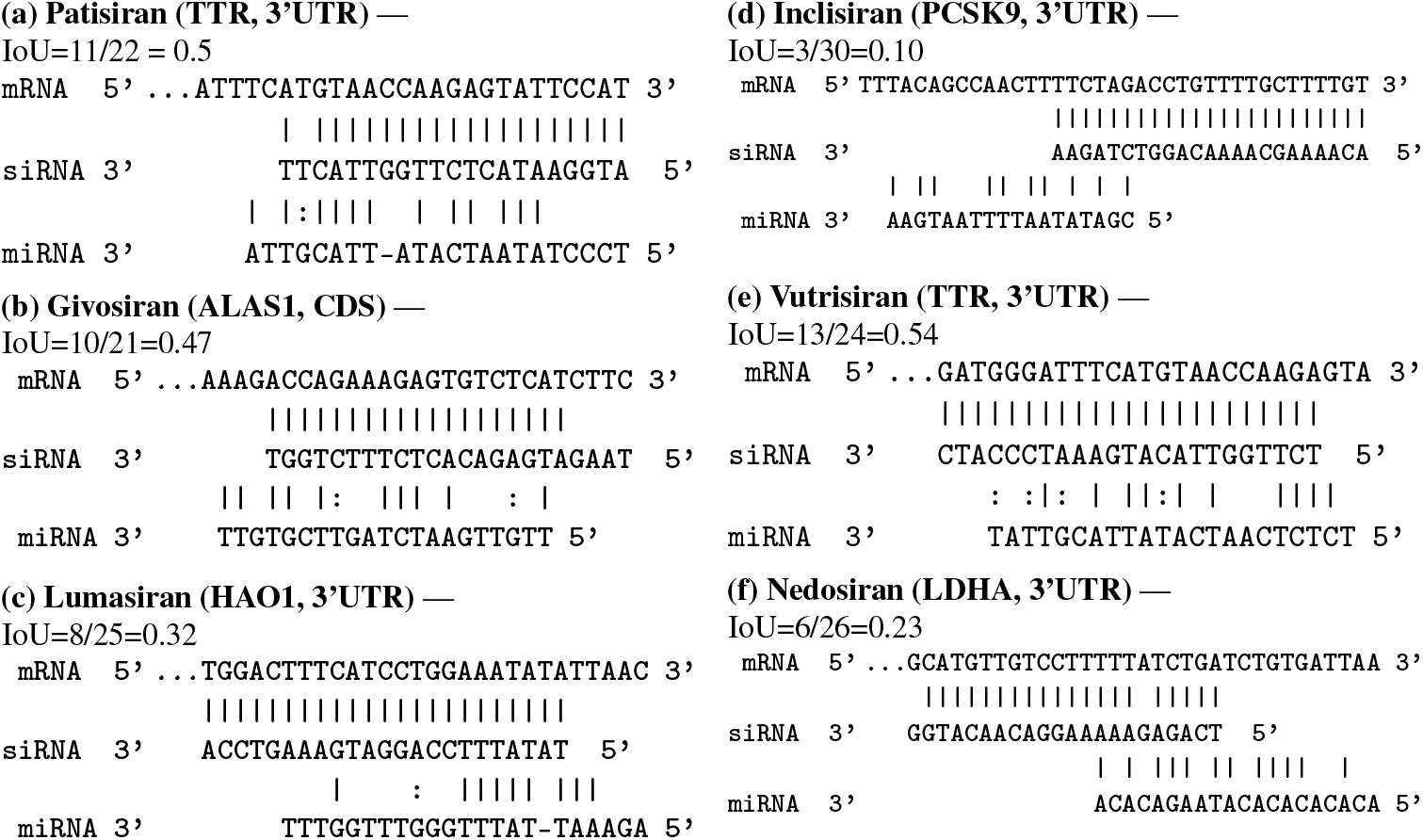
Smith–Waterman local alignments of SpeciMiR-generated miRNAs to target mRNA regions of FDA-approved siRNA drugs. (|): Watson–Crick pairs. (:): G:U wobble pairs. Spaces: mismatches. IoU: Intersection over Union = |*P*_miRNA *∩* siRNA_| / |*P*_miRNA*∪* siRNA_| where *P* is either Watson-Crick or wobble base pairs. The generator was trained on miRNA–mRNA interactions and has never seen siRNA data.

We also scored the thermodynamics of the generated miRNA by measuring change in co-folding energy by RNACofold and MFE by RNAHybrid (Table A.5). We have observed that the generated miRNAs achieved at most 55% of the MFE produced by Nedosiran, and 53% of MFE produced by Patisiran. This energy difference shows the difference between siRNA and miRNA binding mechanism - the FDA-approved siRNA binds to target mRNAs with almost perfect Watson-Crick base pairs, while the SpeciMiR-generated miRNA function through partial binding to the target sites, which has been observed in earlier studies [5].

We further evaluated the generated miRNA and siRNAs on specificity measurements. For each siRNA target mRNA, we identified the siRNA seed and took its reverse complement to find binding site in mRNAs. Then we retrieved 1000 off-target mRNAs from the Manakov2022 test set that share the same seeds as the siRNA. Then we scored the siRNA ↔ off-targets and SpeciMiR-generated miRNA ↔ off-targets with our trained discriminator, RNACofold and RNAhybrid (Table A.5). The SpeciMiR-generated miRNAs exhibit higher specificity scores than the siRNAs (Mean specificity ratio 1.89 v.s. 1.59). This is expected because the discriminator is trained on only miRNA-mRNA pairs, it prefers to score natural miRNA-mRNA higher than unseen siRNA-mRNA interactions. Both RNACofold [20] and RNAhybrid [21] show that siRNAs have higher specificity than the generated miRNAs (Mean specificity ratio 1.60 v.s. 3.66, 1.42 v.s. 2.34). Nevertheless, we observed that the SpeciMiR-generated miRNAs produced higher energies with off-target mRNAs than with on-target mRNAs. Therefore, the generated miRNAs possess structural specificity that binds on-target mRNAs stronger than off-target mRNAs, despite the absence of structural information in the training data.

One outlier is Lumasiran to which we observed higher energy on off-target mRNAs than on-target mRNAs for the SpeciMiR-generated miRNAs. Notably, the synthetic miRNA also has the highest discriminator score on off-target mRNAs (0.557) and the IoU to Lumasiran target mRNA is only 0.32. We speculate that the cause might be due to the non-specific binding to off-target mRNAs that have stronger complementarity with the generated miRNA.

## 5 Discussion

In this study, we developed SpeciMiR, a target-conditioned framework for synthesizing miRNA-like RNAi molecules with improved target specificity. SpeciMiR consists of a mRNA encoder and a Transformer decoder to generate miRNA. The proposed training loss uses a specificity-guided objective, where a discriminator distinguishes true miRNA–mRNA interactions from seed-matched negatives. The generated miRNAs contain canonical target-complementary seed regions and show higher predicted specificity for on-target mRNAs than for seed-matched off-targets. As a proof-of-concept study, SpeciMiR-generated miRNAs bind to similar regions of the target mRNAs for the disease genes as the FDA-approved siRNA drugs for three of six targets and produced candidates with stronger predicted structural compatibility to on-target than off-target mRNAs. Several limitations remain. First, the experimental eCLIP/CLASH training data are limited to 50-nt mRNA contexts, which may restrict generalization to full-length 3’UTRs. Training SpeciMiR on longer target contexts, enabled by sliding-window cross-attention [16], may improve transcript-scale RNAi design. Second, SpeciMiR currently models mRNA-miRNA interactions without explicitly considering RNA secondary structure, RNA-binding proteins such as AGO2 and Dicer. Because miRNAs and siRNAs act through RNA-induced silencing complex (RISC), future work should incorporate RNA– protein co-folding, protein-aware representations, or structure-guided objectives to better capture realistic RNAi activity. Moreover, future model will need to take into account transcriptome context in the host when designing exogenous siRNA. With advance of the RNA sequencing technologies, it is possible to design a cocktail of multiple siRNAs to target specific cell type or pathways. Together, SpeciMiR is an important step towards AI-accelerated RNAi therapeutic design.

## A Appendix

### A.1 SpeciMiR training details

#### Hyperparameter tuning

We optimized the model’s hyperparameters to identify the configuration that yielded the best overall performance. The set of hyperparameters that strike a balance between minimizing validation loss, binding accuracy and F1 score for overlap between the predicted seed match and the TargetScan seed match on the validation set were listed in Table A.2.

#### Decoder training and discriminator finetuning

We pre-trained the decoder on TargetScan dataset [19] to generate miRNA by conditioning on a given mRNA sequence. We trained the decoder on all three mRNA sequence lengths: 30nt, 100nt and 500nt and show the performance in Results4. We trained the discriminator on Manakov2022 training set for 15 epochs. Then we frozen the trained discriminator and used it to guide training of decoder on miRNA synthesis. To finetuned the decoder on specificity loss-guided miRNA synthesis, we trained the decoder on training set for 9 epochs and evaluated our model on validation set.

#### Decoder training

We trained decoder in two stages: 1. Pre-training: we trained the decoder by conditioning miRNA synthesis on 30nt, 100nt, and 500nt mRNA sequences containing seed matches (Experiments 3.1). We trained 2. Finetuning: we finetuned the decoder on Manakov2022 training set for 9 epochs to generalize better to experimental eCLIP/CLASH data. Hyperparameters are selected based on optimized loss on validation sets.

#### Compute Resources

All experiments were conducted on a single NVIDIA RTX 6000 (24 GB) with 32 GB system memory. Phase 1 discriminator training completed in approximately 10 hours; Phase 2 generator training completed in approximately 24 hours.

### A.2 Related Works

#### Enumerate-and-rank pipelines for small RNA design

Computational design of artificial miRNAs and siRNAs enumerates and score each candidate, and filter for specificity post-hoc. WMD3 [10] enumerates 21-mer candidates and filters by thermodynamic asymmetry rules and genome-wide off-target searches for plant amiRNA design. P-SAMS [11] extends this with k-mer-based off-target filtering. miR-Synth [24] adapts the approach for multi-target synthetic miRNAs using regression-based scoring of seed type, structural accessibility, and flanking AU content. More recent tools introduce learned scoring functions while retaining the same paradigm. miRarchitect [23] combines a feedforward neural network for guide strand scoring with multi-stage off-target filtering and scaffold selection from over 16,000 human pri-miRNA backbones. OligoFormer [12] chains convolutional, BiLSTM, and Transformer encoder layers with RNA foundation model embeddings to predict siRNA efficacy, followed by separate off-target assessment via PITA and TargetScan context scores. These ML-guided tools improve the *scoring* step but decouple the specificity from the *generation* step.

#### Generative models for RNA sequences

Recent work has demonstrated that deep generative models can design functional RNA molecules. GEMORNA [15] uses a Transformer encoder-decoder to design mRNA coding sequences optimized for translational efficiency and stability. EVA [14] is a 1.4B-parameter decoder-only Transformer with mixture-of-experts backbone, trained with dual causal and generalized language modeling objectives across 11 RNA classes including miRNA. Although EVA can generate human miRNA sequences conditioned on an RNA-type tag, it only generates from a family-level prior without target mRNA conditioning — it cannot design a miRNA that specifically targets a given mRNA.

#### Present work

SpeciMiR bridges these lines of work by introducing the first end-to-end generative framework for target-conditional miRNA design with integrated specificity optimization. Unlike enumerate-and-rank pipelines, SpeciMiR generates miRNA sequences de novo using a Transformer encoder-decoder conditioned on any target mRNA sequence. Unlike unconditional RNA generators such as EVA, SpeciMiR optimizes for target-specific binding through a frozen discriminator that provides differentiable specificity gradients via a soft miRNA embedding, jointly maximizing on-target binding and minimizing off-target interactions during training.

### A.3 Additional experiments

#### Input-length ablation

We trained SpeciMiR decoders conditioned on 30-, 100-, and 500-nt mRNA inputs using the TargetScan dataset(Section 3.1). All models were trained for 15 epochs on a single GPU using the same optimization protocol. We reported per-token generation accuracy for each input length to assess the effect of conditioning-context length.

#### Three Scorers measure binding affinities between the generated miRNAs and target mRNAs

We scores the binding affinities using three scorers: our finetuned discriminator measuring binding probabilities, RNACofold [20] measuring change in energy in cofolding two RNA sequences (Δ*G*) and RNAHybrid [21] measuring the minimum free energy (MFE) of aligning RNA sequences. We constructed four datasets: real miRNAs ↔ target mRNAs, generated miRNAs ↔ target mRNAs, generated miRNAs ↔ off-target mRNAs, and random miRNAs ↔ target mRNAs. To ensure reasonable running time and sufficient statistics power, we sampled 2000 miRNA-mRNA pairs from Manakov2022 test sets [18] to be real miRNAs ↔ target mRNAs, and generated miRNA for each target mRNA to build a set of generated miRNAs target ↔ mRNAs pairs. For every target mRNA, 5 off-target mRNAs are sampled from all mRNAs targets excluding the target mRNA itself, resulting in 10,000 off-target mRNAs. Random miRNAs are created to match the length of the true miRNAs but they do not contain binding sites. Then we measured the binding probability, Δ*G* and MFE between each miRNA-mRNA pair and measured P-values between each two distributions of scores. All P-values are corrected by sample sizes (Fig.A.3).

#### Benchmarking with miRNA design methods

For miRarchitect, we manually uploaded 40 target sequences from Manakov2022 test set to https://mirarchitect.cs.put.poznan.pl/ and generated miRNA candidates. We picked the one with the highest specificity score to be the best candidate for the given target sequence. For the targets to which no miRNA passes miRarchitect’s filter, we excluded those targets. Overall, 19 miRNAs were generated out of 40 targets. For EVA, we specified the RNA type to be miRNA and species to be human and asked EVA to generate 1000 miRNA candidates with max length of 24nt. Then for each mRNA target, we randomly subsample 50 candidates and rank them by mirna-mrna binding MFE measured by RNAhybrid [21]. The mirna that has the best MFE was picked to be the best candidate. To benchmark specificity of the generated miRNAs, we sampled 20 off-target mRNAs that share the same seed region as the real miRNAs for each target mRNA. We scored the interactions between the generated miRNAs with target mRNAs, and with off-target mRNAs on their binding probability by discriminator *D*_*ϕ*_, Δ_*G*_ by RNACofold [20] and MFE RNAHybrid [21] (Table A.7).

#### siRNA case study

We extracted FDA-approved siRNA sequences from Trader and Yu [5], and collected full target mRNA sequences from NCBI using accession numbers and gene names which are provided by [5] and listed in Table A.8.

### A.4 Supplementary figures

**Figure A.1:**
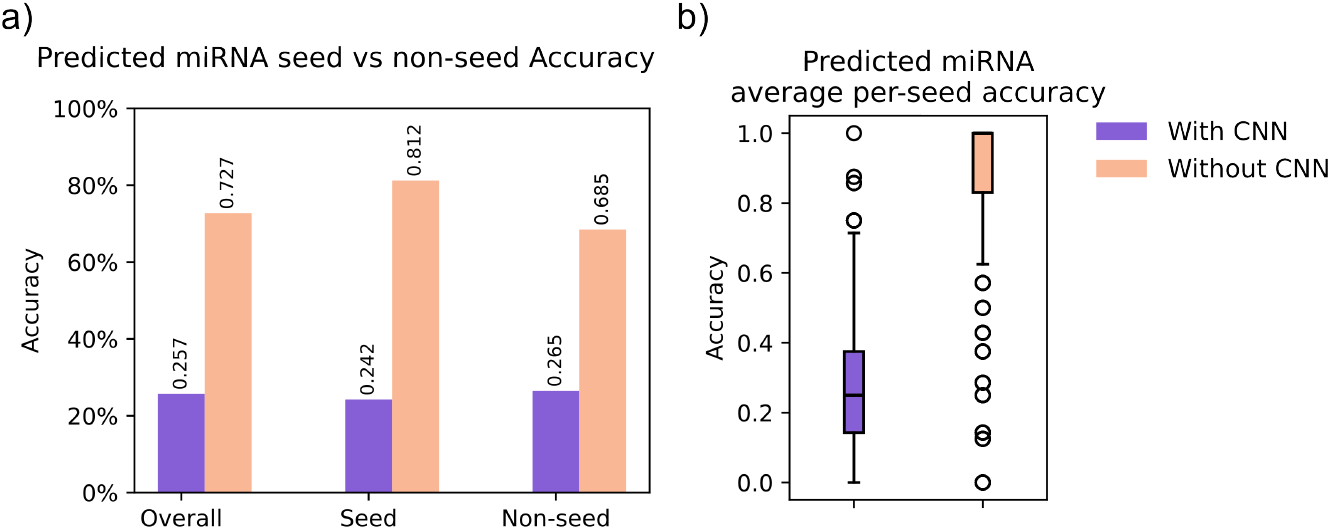
Comparison of generated miRNA accuracy between MiRformer with and without convolution. a) Comparison of predicted miRNA per-base accuracy overall regions, seed and non-seed regions between MiRformer with and without CNN. b) Comparison of distribution of predicted miRNA per-base accuracy in the seed regions between MiRformer with and without convolution.

**Figure A.2:**
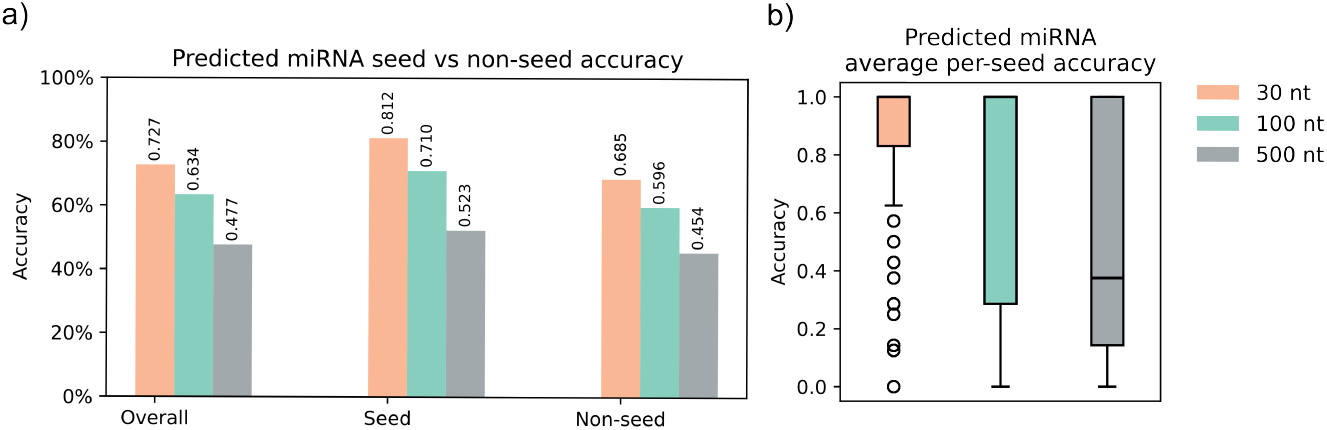
Comparison of generated miRNA accuracy between 30nt, 100nt, 500nt mRNA lengths. a) Comparison of predicted miRNA per-base accuracy overall regions, seed and non-seed regions between 30nt, 100nt and 500nt mRNAs. b) Comparison of distribution of predicted miRNA per-base accuracy in the seed regions between 30nt, 100nt and 500nt mRNAs.

**Figure A.3:**
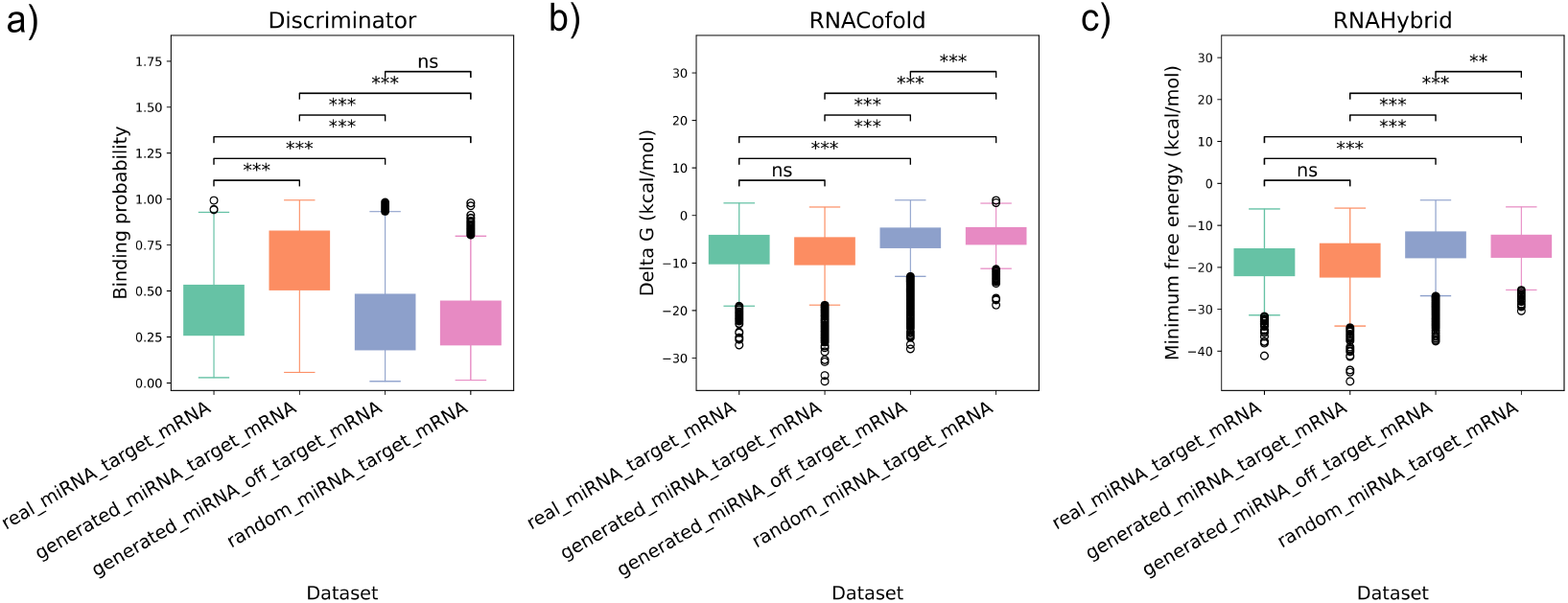
All three scorers on all four datasets show strong binding between the generated miRNAs and target mRNAs. Astricks indicate P-value significance level - ‘*’ is P-value ≤ 0.05; ‘**’ is P-value ≤ 0.01; ‘***’ is P-value ≤ 0.001. ‘ns’ means not significant or P-value ≥ 0.05. All scores are measured on 2000 samples, and 10000 for off-target with K=5 for each miRNA. a) Discriminator-predicted score distribution. Higher scores mean stronger binding. b) RNACofold-predicted score distribution. Lower scores mean stronger binding. c) RNAHybrid-predicted score distribution. Lower scores mean stronger binding.

### A.5 Supplementary tables

**Table A.1:** Symbol notations summary.

| Symbol | Description |
| --- | --- |
| $\theta$ | Generator (decoder) parameters |
| $y_t$ | True miRNA token at position $t$ |
| $T$ | miRNA sequence length |
| $\mathbf{x}_{\text{target}}$ | Target mRNA sequence |
| $\mathbf{x}_{\text{neg}}^{(k)}$ | $k$-th negative (off-target) mRNA sequence |
| $K$ | Number of negative mRNAs per sample |
| $\mathbf{z}_t \in \mathbb{R}^V$ | Generator logits at position $t$ |
| $\mathbf{E}_D \in \mathbb{R}^{V \times D}$ | Frozen discriminator embedding matrix |
| $\tilde{\mathbf{e}}_t \in \mathbb{R}^D$ | Soft embedding at position $t$ |
| $D_\phi(\tilde{\mathbf{e}}, \mathbf{x})$ | Discriminator interaction probability |
| $\eta_{\text{slow}} = 5\text{e-}5$ | Learning rate for cross attention layer |
| $\eta_{\text{fast}} = 1\text{e-}4$ | Learning rate for discriminator head |
| $\tau = 0.5$ | Softmax temperature |
| $\alpha = 0.5, \beta = 0.7$ | Loss weights for generation and specificity |
| $\lambda = 0.5$ | Off-target penalty weight |

**Table A.2:**
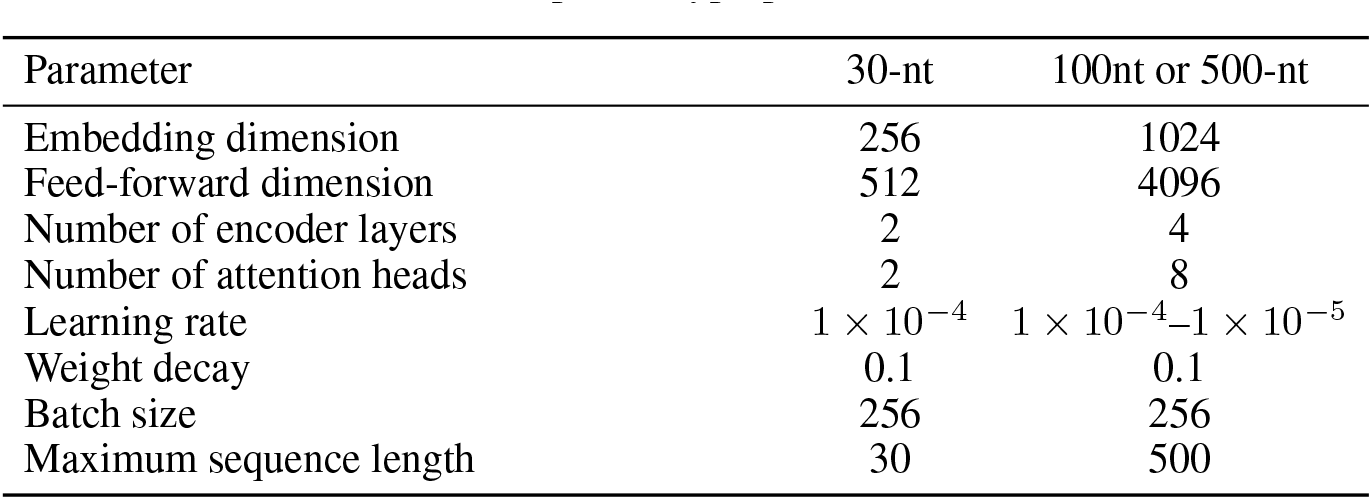
Optimal hyperparameters for SpeciMiR model.

| Parameter | 30-nt | 100nt or 500-nt |
| --- | --- | --- |
| Embedding dimension | 256 | 1024 |
| Feed-forward dimension | 512 | 4096 |
| Number of encoder layers | 2 | 4 |
| Number of attention heads | 2 | 8 |
| Learning rate | $1 \times 10^{-4}$ | $1 \times 10^{-4}$–$1 \times 10^{-5}$ |
| Weight decay | 0.1 | 0.1 |
| Batch size | 256 | 256 |
| Maximum sequence length | 30 | 500 |

**Table A.3:** Comparison of predicted miRNA seed match types across different mRNA lengths.

| Metric | 30nt | 100nt | 500nt |
| --- | --- | --- | --- |
| *Raw Counts* |  |  |  |
| Valid 6-mers | 48 | 17 | 8 |
| Valid 7-mers-A1 | 902 | 939 | 844 |
| Valid 7-mers-m8 | 2,180 | 2,244 | 2,240 |
| Valid 8-mers | 2,249 | 2,495 | 2,580 |
| **Total Functional Match Rate** | **94.17%** | **99.70%** | **99.30%** |
| *Distribution (% of Total Matches)* |  |  |  |
| Percentage of 6-mers | 0.89% | 0.30% | 0.14% |
| Percentage of 7-mers-A1 | 16.77% | 16.49% | 14.88% |
| Percentage of 7-mers-m8 | 40.53% | 39.40% | 39.49% |
| Percentage of 8-mers | 41.81% | 43.81% | 45.49% |

**Table A.4:** Seed match verification on perturbed mRNA sequences across mRNA context lengths. Each model generates an miRNA from the perturbed mRNA; a match indicates the generated miRNA still finds a seed site in the perturbed mRNA sequence.

| **Metric** | **30nt** | **100nt** | **500nt** |
| --- | --- | --- | --- |
| *Raw Counts* |  |  |  |
| Valid 6-mers | 372 | 189 | 156 |
| Valid 7-mers-A1 | 540 | 1,252 | 1,105 |
| Valid 7-mers-m8 | 532 | 1,946 | 2,214 |
| Valid 8-mers | 301 | 1,396 | 1,672 |
| **Total Seed Match Rate** | **30.55%** | **83.74%** | **90.11%** |
| *Distribution (% of Total Matches)* |  |  |  |
| Percentage of 6-mers | 21.32% | 3.95% | 3.03% |
| Percentage of 7-mers-A1 | 30.95% | 26.18% | 21.47% |
| Percentage of 7-mers-m8 | 30.49% | 40.69% | 43.02% |
| Percentage of 8-mers | 17.25% | 29.19% | 32.48% |

**Table A.5:** Seed match verification on perturbed mRNA sequences across mRNA context lengths using the entire miRNA. Each model generates an miRNA from the perturbed mRNA; a match indicates the generated miRNA finds a complementary site anywhere along its length in the perturbed mRNA.

| **Metric** | **30nt** | **100nt** | **500nt** |
| --- | --- | --- | --- |
| *Raw Counts* |  |  |  |
| Valid 6-mers | 648 | 313 | 181 |
| Valid 7-mers-A1 | 366 | 657 | 435 |
| Valid 7-mers-m8 | 817 | 2,182 | 2,340 |
| Valid 8-mers | 559 | 2,031 | 2,639 |
| **Total Seed Match Rate** | **41.84%** | **90.74%** | **97.95%** |
| *Distribution (% of Total Matches)* |  |  |  |
| Percentage of 6-mers | 27.11% | 6.04% | 3.24% |
| Percentage of 7-mers-A1 | 15.31% | 12.68% | 7.77% |
| Percentage of 7-mers-m8 | 34.18% | 42.10% | 41.82% |
| Percentage of 8-mers | 23.39% | 39.19% | 47.17% |

**Table A.6:** Specificity-guided generator training and validation metrics. *p*_*on*_=binding probability on-target mRNAs. *p*_*off*_ =binding probability on off-target mRNAs.

| **Metric** | **Train** | **Val** |
| --- | --- | --- |
| Total Loss ($\alpha\mathcal{L}_{gen} + \beta\mathcal{L}_{spec}$) | 0.4730 | 0.4664 |
| Generation Loss ($\mathcal{L}_{gen}$) | 0.3206 | 0.3183 |
| Specificity Loss ($\mathcal{L}_{spec}$) | 0.4467 | 0.4389 |
| On-target binding probability ($p_{on}$) | 0.845 | 0.852 |
| Off-target binding probability ($p_{off}$) | 0.335 | 0.335 |
| Specificity Gap ($p_{on} - p_{off}$) | 0.510 | 0.517 |

**Table A.7:** Comparison with miRNA designing methods. Scores are reported as mean ± std. **Converage** proportion of mRNA target sequences receiving a valid miRNA design. **Specificity gap** (|on−off|, ↑) measures gap between on and off targets; ↑ means larger the gap the better.

| Method | Condition | $D_\phi$<br>($p_{\text{bind}}$) | RNAcofold<br>($\Delta_G$) | RNAhybrid<br>(MFE) | Target-specific<br>generation | Sequence space<br>exploration |
| --- | --- | --- | --- | --- | --- | --- |
| *Reference* |  |  |  |  |  |  |
| | Observed miRNA ↔ target | $0.42 \pm 0.2$ | $-7.2 \pm 4.5$ | $-19.0 \pm 4.5$ | | |
| **SPECIMiR (ours)** |  |  |  |  | ✓ | ✓ |
| | Generated ↔ target | $0.70 \pm 0.2$ | $-8.0 \pm 4.0$ | $-18.0 \pm 4.0$ | | |
| | Generated ↔ off-target | $0.40 \pm 0.2$ | $-5.4 \pm 4.0$ | $-15.0 \pm 4.0$ | | |
| | Specificity gap (↑) | **0.30** | 3.4 | 3.1 | | |
| **miRarchitect [23]** |  |  |  |  | ✓ | ✗ |
| | Generated ↔ target | $0.41 \pm 0.2$ | $-28.2 \pm 3.1$ | $-41.0 \pm 3.1$ | | |
| | Generated ↔ off-target | $0.30 \pm 0.2$ | $-5.0 \pm 3.1$ | $-14.6 \pm 3.5$ | | |
| | Specificity gap (↑) | 0.11 | **23.2** | **26.4** | | |
| **EVA [14]<sup>1</sup>** |  |  |  |  | ✗ | ✓ |
| | Generated ↔ target | $0.22 \pm 0.2$ | $-3.4 \pm 2.2$ | $-13.7 \pm 4.4$ | | |
| | Generated ↔ off-target | $0.26 \pm 0.2$ | $-4.1 \pm 2.7$ | $-13.9 \pm 4.1$ | | |
| | Specificity gap (↑) | 0.04 | 0.7 | 0.2 | | |

<sup>1</sup> EVA is unconditional on target mRNA sequences. We provided the comparison for general generation quality benchmark.

**Table A.8:**
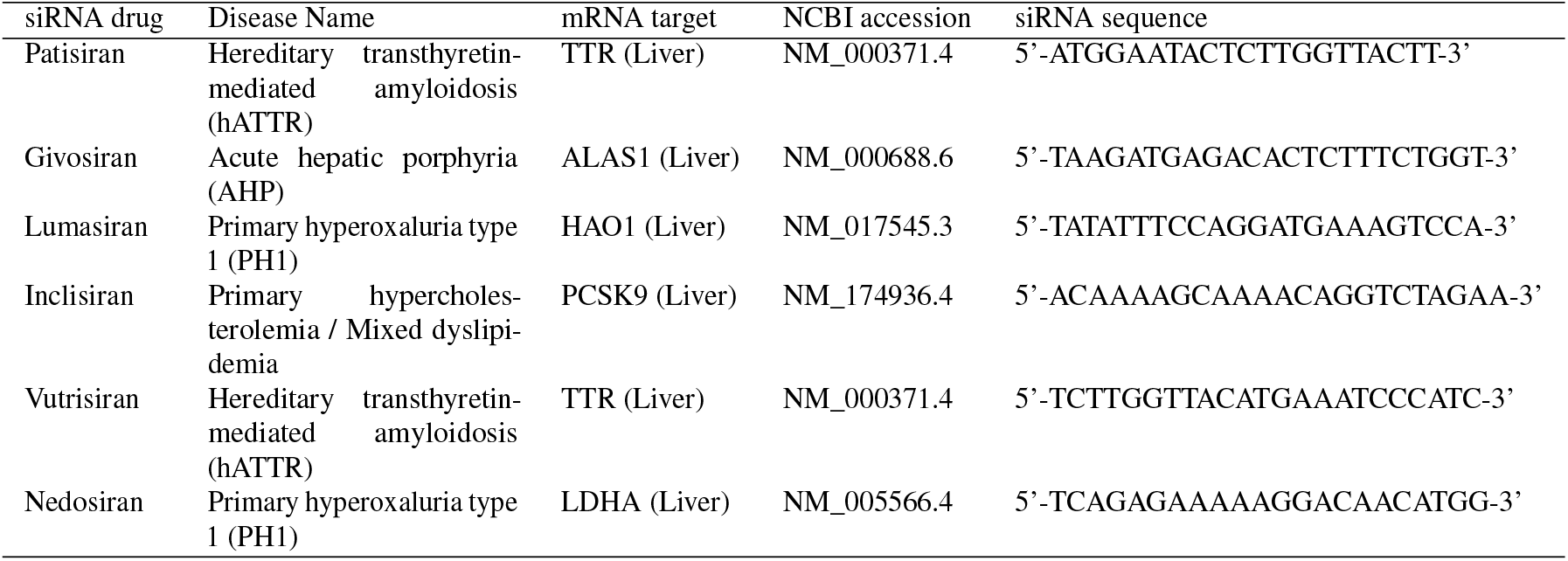
FDA-approved siRNA sequences and mRNA target sources.

**Table A.9:** Thermodynamic comparison between SpeciMiR-generated miRNAs and FDA-approved siRNA guide strands against the same target mRNA regions. RNACofold: co-folding Δ*G*_binding_ (kcal/mol). RNAhybrid: minimum free energy of best binding site (kcal/mol). More negative values indicate stronger binding. Energy ratio = |Δ*G*_generated_|*/*|Δ*G*_FDA_| or |MFE_generated_|*/*|MFE_FDA_|

| Drug | Target | RNAcofold $\Delta G_{\text{bind}}$ (kcal/mol) | | | RNAhybrid MFE (kcal/mol) | | |
| --- | --- | --- | --- | --- | --- | --- | --- |
|  |  | FDA | Ours | Ratio | FDA | Ours | Ratio |
| Patisiran | TTR | -20.44 | -5.32 | 26.0% | -35.1 | -18.5 | 52.7% |
| Givosiran | ALAS1 | -31.55 | -4.92 | 15.6% | -43.1 | -13.0 | 30.2% |
| Lumasiran | HAO1 | -30.02 | -4.63 | 15.4% | -42.6 | -13.0 | 30.5% |
| Inclisiran | PCSK9 | -33.03 | -6.69 | 20.3% | -40.6 | -12.7 | 31.3% |
| Vutrisiran | TTR | -28.31 | -3.22 | 11.4% | -43.4 | -17.3 | 39.9% |
| Nedosiran | LDHA | -22.39 | -8.09 | **36.1%** | -33.5 | -18.3 | **54.6%** |

**Table A.10:** On-target vs. off-target specificity comparison between SpeciMiR-generated miRNAs and FDA-approved siRNAs. Off-targets are seed-matched mRNAs from the Manakov2022 test set sharing the same 7mer seed site as the target. *D*_*ϕ*_: discriminator score (higher = stronger binding). RNACofold Δ*G*_bind_ and RNAhybrid MFE in kcal/mol (more negative = stronger binding). Specificity Ratio=On/Off where a larger ratio = more specific.

| Drug | Src | $D_\phi$ | | | RNACofold (kcal/mol) | | | RNAhybrid (kcal/mol) | | |
| --- | --- | --- | --- | --- | --- | --- | --- | --- | --- | --- |
|  |  | On | Off | Specificity | On | Off | Specificity | On | Off | Specificity |
|  |  |  |  | Ratio |  |  | Ratio |  |  | Ratio |
| Patisiran | Ours | 0.771 | 0.388 | **1.99** | -5.32 | -2.73 | 1.95 | -18.5 | -10.01 | 1.85 |
|  | FDA | 0.796 | 0.408 | 1.95 | -20.4 | -7.85 | **2.60** | -35.1 | -16.55 | **2.12** |
| Givosiran | Ours | 0.595 | 0.352 | 1.69 | -4.92 | -3.91 | 1.26 | -13.01 | -12.5 | 1.05 |
|  | FDA | 0.628 | 0.341 | **1.84** | -31.6 | -6.33 | **4.99** | -43.1 | -16.8 | **2.57** |
| Lumasiran | Ours | 0.811 | 0.557 | **1.46** | -4.63 | -4.51 | 1.03 | -13.0 | -13.8 | 0.95 |
|  | FDA | 0.457 | 0.388 | 1.18 | -30.0 | -5.01 | **5.99** | -42.6 | -15.5 | **2.75** |
| Inclisiran | Ours | 0.664 | 0.369 | **1.80** | -6.69 | -2.53 | 2.64 | -12.7 | -9.09 | 1.40 |
|  | FDA | 0.475 | 0.398 | 1.19 | -33.0 | -8.64 | **3.82** | -40.6 | -18.1 | **2.24** |
| Vutrisiran | Ours | 0.758 | 0.377 | **2.01** | -3.22 | -3.18 | 1.01 | -17.3 | -10.1 | 1.71 |
|  | FDA | 0.848 | 0.430 | 1.97 | -28.3 | -7.53 | **3.76** | -43.4 | -17.0 | **2.55** |
| Nedosiran | Ours | 0.890 | 0.324 | **2.75** | -8.09 | -3.54 | 2.29 | -18.3 | -10.1 | 1.81 |
|  | FDA | 0.486 | 0.361 | 1.35 | -22.4 | -9.59 | **2.34** | -33.5 | -18.1 | **1.85** |
| Mean | Ours | 0.748 | 0.395 | **1.89** | -5.48 | -3.43 | 1.60 | -15.5 | -10.9 | 1.42 |
|  | FDA | 0.615 | 0.388 | 1.59 | -27.62 | -7.54 | **3.66** | -39.7 | -17.0 | **2.34** |

## References

[1] David P Bartel. Micrornas: genomics, biogenesis, mechanism, and function. cell, 116(2):281– 297, 2004.

[2] Bo Wang, Shu-hao Hsu, Xinmei Wang, Huban Kutay, Hemant Kumar Bid, Jianhua Yu, Ramesh K Ganju, Samson T Jacob, Mariia Yuneva, and Kalpana Ghoshal. Reciprocal regulation of microrna-122 and c-myc in hepatocellular cancer: role of e2f1 and transcription factor dimerization partner 2. Hepatology, 59(2):555–566, 2014.

[3] Han Han, Dan Sun, Wenjuan Li, Hongxing Shen, Yahui Zhu, Chen Li, Yuxing Chen, Longfeng Lu, Wenhua Li, Jinxiang Zhang, et al. A c-myc-microrna functional feedback loop affects hepatocarcinogenesis. Hepatology, 57(6):2378–2389, 2013.

[4] Yiran Zhu, Liyuan Zhu, Xian Wang, and Hongchuan Jin. RNA-based therapeutics: an overview and prospectus. 13(7):644, 2022.

[5] Gavin M. Traber and Ai-Ming Yu. The growing class of novel RNAi therapeutics. 106(1):13–20, 2024.

[6] Aimee L. Jackson and Peter S. Linsley. Recognizing and avoiding siRNA off-target effects for target identification and therapeutic application. 9(1):57–67, 2010.

[7] Julia Neumeier and Gunter Meister. siRNA specificity: RNAi mechanisms and strategies to reduce off-target effects. Volume 11-2020.

[8] Dieter Huesken, Joerg Lange, Craig Mickanin, Jan Weiler, Fred Asselbergs, Justin Warner, Brian Meloon, Sharon Engel, Avi Rosenberg, Dalia Cohen, Mark Labow, Mischa Reinhardt, François Natt, and Jonathan Hall. Design of a genome-wide siRNA library using an artificial neural network. 23(8):995–1001, 2005.

[9] Bowen Xiao, Shaopeng Wang, Yu Pan, Wenjun Zhi, Chensheng Gu, Tao Guo, Jiaqi Zhai, Chenxu Li, Yong Q. Chen, and Rong Wang. Development, opportunities, and challenges of siRNA nucleic acid drugs. 36(1), 2025.

[10] Rebecca Schwab, Stephan Ossowski, Markus Riester, Norman Warthmann, and Detlef Weigel. Highly specific gene silencing by artificial MicroRNAs in arabidopsis. 2026.

[11] Noah Fahlgren, Steven T Hill, James C Carrington, and Alberto Carbonell. P-sams: a web site for plant artificial microrna and synthetic trans-acting small interfering rna design. Bioinformatics, 32(1):157–158, 2016.

[12] Yilan Bai, Haochen Zhong, Taiwei Wang, and Zhi John Lu. Oligoformer: an accurate and robust prediction method for sirna design. Bioinformatics, 40(10):btae577, 2024.

[13] Sergei A Manakov, Alexander A Shishkin, Brian A Yee, Kylie A Shen, Diana C Cox, Samuel S Park, Heather M Foster, Karen B Chapman, Gene W Yeo, and Eric L Van Nostrand. Scalable and deep profiling of mrna targets for individual micrornas with chimeric eclip. BioRxiv, pages 2022–02, 2022.

[14] Yanjie Huang, Guangye Lv, Anyue Cheng, Wei Xie, Mengyan Chen, Xinyi Ma, Yijun Huang, Yueyang Tang, Qingya Shi, Jiahao Wang, et al. A long-context generative foundation model deciphers rna design principles. bioRxiv, pages 2026–03, 2026.

[15] He Zhang, Hailong Liu, Yushan Xu, Haoran Huang, Yiming Liu, Jia Wang, Yan Qin, Haiyan Wang, Lili Ma, Zhiyuan Xun, et al. Deep generative models design mrna sequences with enhanced translational capacity and stability. Science, 390(6773):eadr8470, 2025.

[16] Jiayao Gu, Can Chen, and Yue Li. Mirformer: a dual-transformer-encoder framework for predicting microrna-mrna interactions from paired sequences. bioRxiv, 2025.

[17] Iz Beltagy, Matthew E. Peters, and Arman Cohan. Longformer: The long-document transformer. (2004.05150), 2020.

[18] Stephanie Sammut, Katarina Gresova, Dimosthenis Tzimotoudis, Eva Marsalkova, David Cechak, and Panagiotis Alexiou. miRBench: novel benchmark datasets for microrna binding site prediction that mitigate against prevalent microrna frequency class bias. Bioinformatics, 41(Supplement_1):i542–i551, 2025.

[19] Vikram Agarwal, George W Bell, Jin-Wu Nam, and David P Bartel. Predicting effective microrna target sites in mammalian mrnas. elife, 4:e05005, 2015.

[20] Ronny Lorenz, Stephan H. Bernhart, Christian Höner zu Siederdissen, Hakim Tafer, Christoph Flamm, Peter F. Stadler, and Ivo L. Hofacker. ViennaRNA package 2.0. 6(1):26, 2011.

[21] Jan Krüger and Marc Rehmsmeier. RNAhybrid: microRNA target prediction easy, fast and flexible. 34:W451–W454, 2006.

[22] David P Bartel. Metazoan micrornas. Cell, 173(1):20–51, 2018.

[23] Agnieszka Belter, Jaroslaw Synak, Marta Mackowiak, Anna Kotowska-Zimmer, Marek Figlerowicz, Marta Szachniuk, and Marta Olejniczak. Machine learning-guided design of artificial microRNAs for targeted gene silencing. 2026.

[24] Alessandro Laganà, Mario Acunzo, Giulia Romano, Alfredo Pulvirenti, Dario Veneziano, Luciano Cascione, Rosalba Giugno, Pierluigi Gasparini, Dennis Shasha, Alfredo Ferro, et al. mir-synth: a computational resource for the design of multi-site multi-target synthetic mirnas. Nucleic Acids Research, 42(9):5416–5425, 2014.

